# SSRE: Cell Type Detection Based on Sparse Subspace Representation and Similarity Enhancement

**DOI:** 10.1101/2020.04.08.028779

**Authors:** Zhenlan Liang, Min Li, Ruiqing Zheng, Yu Tian, Xuhua Yan, Jin Chen, Fang-Xiang Wu, Jianxin Wang

## Abstract

Accurate identification of cell types from single-cell RNA sequencing (scRNA-seq) data plays a critical role in a variety of scRNA-seq analysis studies. It corresponds to solving an unsupervised clustering problem, in which the similarity measurement between cells in a high dimensional space affects the result significantly. Although many approaches have been proposed recently, the accuracy of cell type identification still needs to be improved. In this study, we proposed a novel single-cell clustering framework based on similarity learning, called SSRE. In SSRE, we model the relationships between cells based on subspace assumption and generate a sparse representation of the cell-to-cell similarity, which retains the most similar neighbors for each cell. Besides, we adopt classical pairwise similarities incorporated with a gene selection and enhancement strategy to further improve the effectiveness of SSRE. For performance evaluation, we applied SSRE in clustering, visualization, and other exploratory data analysis processes on various scRNA-seq datasets. Experimental results show that SSRE achieves superior performance in most cases compared to several state-of-the-art methods.

## Introduction

With the recent emergence of single-cell RNA sequencing (scRNA-seq) technology, numerous scRNA-seq datasets have been generated, bringing unique challenges for advanced omics data analysis [1,2]. Unlike bulk sequencing averaging the expression of mass cells, scRNA-seq technique quantifies gene expression at the single cell resolution. Single cell techniques promote a wide variety of biological topics such as cell heterogeneity, cell fate decisions and disease pathogenesis [3–5]. Among all the applications, cell type identification plays a fundamental role and its performance has a deep impact on downstream researches [6]. However, identifying cell types from scRNA-seq data is still a challenging problem because of the high noise rate and high dropouts, which cannot be addressed by traditional clustering methods well [7]. Therefore, new efficient and reliable clustering methods for cell type identification are urgent and meaningful.

In recent studies, several novel clustering approaches for detecting cell types from scRNA-seq data have been proposed. Among these methods, cell types are mainly decided on the basis of cell-to-cell similarity learned from scRNA-seq data. SIMLR [8] visualizes and clusters cells using multi-kernel similarity learning [9], which performs well on grouping cells. SNN-Cliq [10] firstly constructs a distance matrix based on the Euclidean distance, and then introduces the shared k-nearest-neighbors model to redefine the similarity. SNN-Cliq provides both the estimation of cluster number and the clustering results by searching for quasi-cliques. Jiang et al [11] proposed the differentiability correlation between pairs of cells instead of computing primary (dis)similarity using the Pearson correlation or the Euclidean distance. RAFSIL [12] divides genes into multiple clusters and concatenates the informative features from each gene cluster after dimension reduction, and finally applies the random forest to calculate the similarities for each cell recursively. Besides, NMF determines the cell types in latent space via nonnegative matrix factorization [13], while SinNLRR [14] learns a similarity matrix with nonnegative and low rank constraints. Instead of learning a specific similarity, some researchers have turned to use ensemble learning based on the consensus of multiple clustering methods in order to obtain robust results [15,16].

Even though many approaches have been applied to cell type identification, most of the previous methods compute the similarity between two cells merely considering their own gene expressions which is sensitive to the noise, especially on data with high dimension [17]. In this study, we develop SSRE, a novel method for cell type identification focused on similarity learning, in which the cell-to-cell similarity is measured by considering more similar neighbors. SSRE computes the linear representation between cells to generate a sparse representation of cell-to-cell similarity based on the sparse subspace theory [18]. Moreover, SSRE incorporates three classical pairwise similarities, motivated by the observations that each similarity measurement can represent data from a different aspect [15,19]. In order to reduce the effect of irrelevant features and to improve the overall accuracy, we design a two-step procedure in SSRE, *i.e.*, 1) adaptive gene selection and 2) similarity enhancement. Experiments show that the new similarities in SSRE, combined with spectral clustering (SC), can reveal the block structure of scRNA-seq data reliably. Also, the experimental results on ten real scRNA-seq datasets and five simulated scRNA-seq datasets show that SSRE achieves higher accuracy on cell type detection in most cases compared with popular approaches. Moreover, we also show that SSRE can be easily extended to other scRNA-seq tasks such as differential expression analysis and data visualization.

## Materials and methods

### Framework of SSRE

We introduce the overview of SSRE briefly. A schematic diagram of SSRE is shown in **Figure 1**, and detailed steps of SSRE will be introduced later in this section. Given a scRNA-seq expression matrix, we first remove genes whose expression are zero in all the cells. Then, the informative genes are selected based on the sparse subspace representation (SSR), Pearson, Spearman and Cosine similarities. With the preprocessed gene expression matrix, SSRE learns sparse representation for each cell simultaneously. Then, SSRE derives an enhanced similarity matrix from these learned sparse similarities. Finally, SSRE uses the enhanced similarity to identify cell types and visualize results.

**Figure 1.**
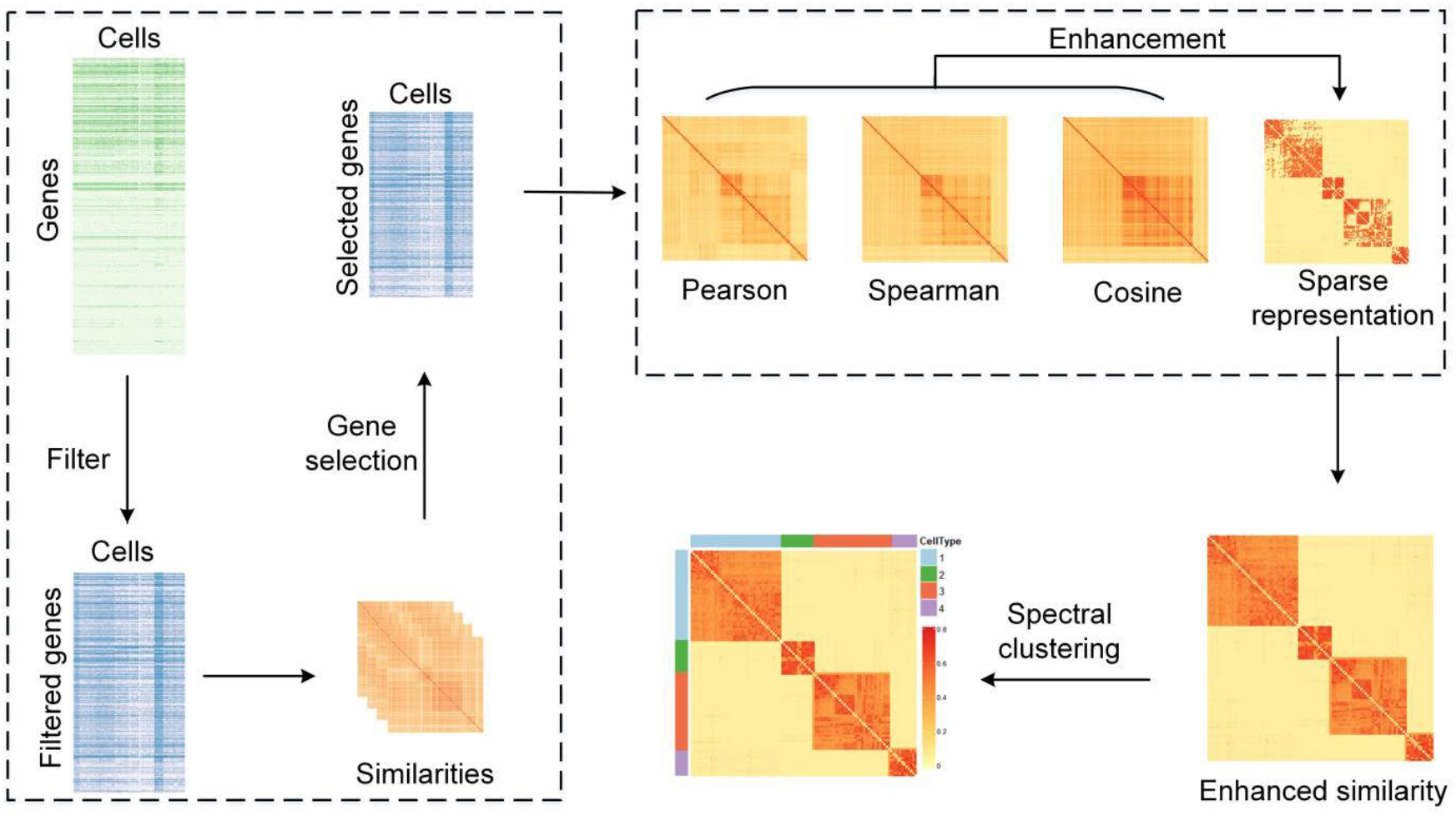
The schematic diagram of SSRE. The main steps of SSRE are displayed, which include gene filtering, gene selection, calculating different similarities, similarity enhancement and clustering.

### Sparse subspace representation

Estimation of the similarity (or distance) matrix is a crucial step in clustering [8]. If the similarity matrix is well generated, it could be relatively easier to distinguish the cluster. In this paper, we adopt sparse subspace theory [18] to compute the linear representation between cells and generate a sparse representation of the cell-to-cell similarity. Some subspace-based clustering methods have been successfully applied to computer vision field and proved to be highly robust in corrupted data [20,21]. For scRNA-seq data, the sparse representation of the cell-to-cell similarity is measured by considering the linear combination of similar neighbors instead of only these two cells, which tends to catch more global structure information and generate more reliable similarity. The specific calculation processes are described as follows.

Mathematically, given a gene expression dataset with *p* genes and *n* cells, denoted by *X* = [*x*_1_, *x*_2_, …, *x*_*n*_] ∈ *R*^*p*×*n*^, where *x*_*i*_ = [*x*_*i*1_, *x*_*i*2_, …, *x*_*ip*_]^*T*^indicates the expression profiles of the *p* genes in cell *i*, the linear representation coefficient matrix *C* = [*c*_1_, *c*_2_, …, *c*_*n*_] ∈ *R*^*n*×*n*^ satisfies the equation *X* = *XC*. With the assumption that the expression of a cell can be represented by the other cells with the same type, only the similarity of cells in the same cluster is non-zero, which means the coefficient matrix *C* is usually sparse. With the relaxed sparse constraint, the coefficient matrix *C* can be computed by solving an optimization problem as follows:

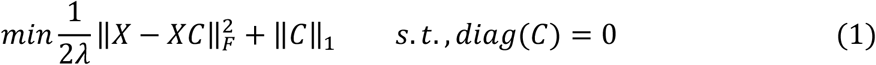

Where ‖ · ‖_*F*_ denotes the Frobenius norm which calculates the square root of sum of all squared elements constraint *diag*(*C*) = 0 prevents the cells from being represented by themselves, while *λ* is a penalty factor. An efficient approach to solve Equation (1) is the alternating direction method of multipliers (ADMM) [22]. We rewrite Equation (1) as follows:

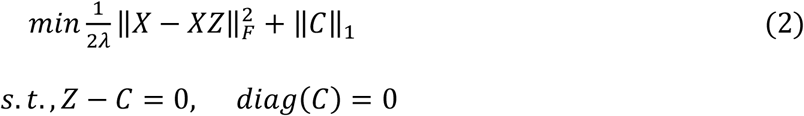

where *Z* is an auxiliary matrix. According to the model of ADMM, the augmented Lagrangian with auxiliary matrix *Z* and penalty parameter (*γ*) > 0 for the optimization formula (2) is

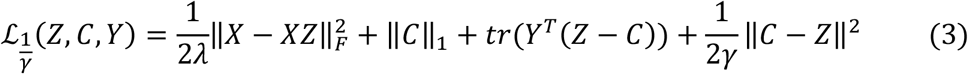

where *Y* is the dual variable. The derivation of its update also can be found in section 1 of File S1. The matrix *C* is the target sparse representation matrix. To keep the symmetry and nonnegative nature of the similarity matrix, the element of sparse representation similarity *sim*_*sparse*_ is calculated as *sim*_*sparse*_(*i*, *j*) = |*c*_*ij*_| + |*c*_*ji*_|. The above similarity learning with sparse constraint is named SSR.

### Data preprocessing and gene selection

Before applying SSR in cell type detection, data preprocessing is required. Various data preprocessing methods have been used in the previous studies, such as gene filter [12,15] and imputation [23,24]. In our method, we first remove genes with zero expression in all of cells and apply *L*_2_-norm to each cell to eliminate the expression scale difference between different cells. Then, we compute the preliminary *sim*_*sparse*_ with the normalized gene expression matrix. Next, we adopt the Laplacian score [25] on *sim*_*sparse*_ to measure the contribution of genes to the learned cell-to-cell similarity and select significant genes for the following study. Genes with higher Laplacian scores are considered as more informative in distinguishing cell types [8]. Besides the sparse similarity *sim*_*sparse*_, we also consider three additional pairwise similarities, *i.e.* Pearson, Spearman, and Cosine, to evaluate the importance of genes (denoted as *sim*_*pearson*_, *sim*_*spearman*_ and *sim*_*cosine*_, respectively). For each similarity, we rank genes in descending order by the Laplacian score and select the top *t* genes as important gene set that is denoted by *G*_1_. The determination of the threshold *t* can be formulated as

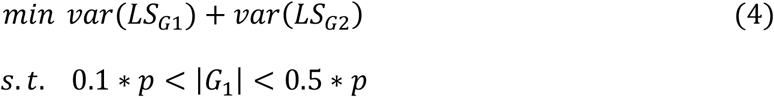

where *G*_1_= [*g*_1_, *g*_2_, … *g*_*t*−1_] and *G*_2_ = [*g*_*t*_, *g*_*t*+1_, … *g*_*p*_] denote two gene sets divided by *t*. The *LS*_*G*1_ and *LS*_*G*2_ are the Laplacian scores of genes in sets *G*_1_ and *G*_2_, respectively, and |∗| is the cardinality of a set. The *var*(∗) indicates variance of a set while *p* is the number of genes. Finally, we recompute *sim*_*sparse*_, *sim*_*pearson*_, *sim*_*spearman*_ and *sim*_*cosine*_ based on the intersection of four selected important gene sets. In the next section, we introduce an enhancement strategy to further improve the learned sparse similarity *sim*_*sparse*_.

### Similarity enhancement

The sparse representation similarity *sim*_*sparse*_ may suffer from the high-level technical noise in the data resulting in underestimation. Inspired by the consensus clustering and resource allocation, we further enhance *sim*_*sparse*_ by integrating multiple pairwise similarities including *sim*_*pearson*_, *sim*_*spearman*_ and *sim*_*cosine*_, which partially reveal the local information between cells.

Based on the similarity matrices described in previous Section, we impute missing values in *sim*_*sparse*_ according to the nearest neighbors’ information in all the three pairwise similarity matrices. We firstly define a target similarity matrix *P* as follows:

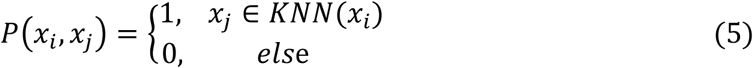

where *KNN*(*x*_*i*_) indicates the k-nearest neighbors of cell *x*_*i*_. Then we mark the similarity *sim*_*sparse*_(*x*_*i*_, *x*_*j*_) between cells *x*_*i*_ and *x*_*j*_ as a missing value when it is zero in the *sim*_*sparse*_ but *P*(*x*_*i*_, *x*_*j*_) = 1 in at least one pairwise similarity matrix. Let *Isim*_*sparse*_ = *O*^*n*×*n*^ denotes the initial matrix to be imputed and *n* is the number of cells. For a marked missing value, the similarity *Isim*_*sparse*_(*x*_*i*_, *x*_*j*_) is computed by the modified Weighted Adamic/Adar [26, 27]. It is formulated as follows:

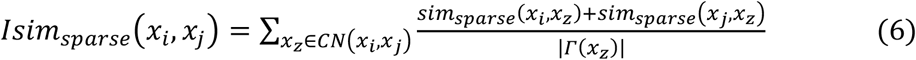

where |*Γ*(*x*_*z*_)| indicates the number of neighbors of cell *x*_*z*_ while *CN*(*x*_*i*_, *x*_*j*_) denotes the set of common neighbors of cell *x*_*i*_ and *x*_*j*_. Note that the imputed similarity *Isim*_*sparse*_(*x*_*i*_, *x*_*j*_) is zero when *CN*(*x*_*i*_, *x*_*j*_) = ∅. At the end of the process, an enhanced and more comprehensive sparse representation matrix *Esim*_*sparse*_ is obtained and computed as *Esim*_*sparse*_ = *Isim*_*sparse*_ + *Isim*_*sparse*_^*T*^+ *sim*_*sparse*_.

### Spectral clustering

Spectral clustering is a typical clustering technique that divides multiple objects into disjoint clusters depending on the spectrum of the similarity matrix [28]. Compared with the traditional clustering algorithms, spectral clustering is advantageous in model simplicity and robustness. In this study, we perform spectral clustering on the final enhanced sparse representation similarity *Esim*_*sparse*_. The inputs of spectral clustering are the cell-to-cell similarity matrix and the cluster number. The detailed introduction and analysis of spectral clustering could be found in previous studies [28,29].

### Datasets

Datasets used in this study consist of two parts, real scRNA-seq dataset and simulated scRNA-seq dataset. The real scRNA-seq datasets are obtained from Gene Expression Omnibus (GEO) [30] and ArrayExpress [31]. We collect ten real scRNA-seq datasets that vary either in terms of species, tissues, and biological processes. They include Treutlein dataset [32], Yan dataset [33], Deng dataset [34], Goolam dataset [35], Ting dataset [36], Song dataset [37], Engel dataset [38], Haber dataset [39], Vento dataset [40], Macosko dataset [41]. The scale of these ten datasets varies from dozens to thousands, and the gene expression values are computed by different units. The details of these real datasets are described in **Table 1**. In addition, we use Splatter [42] to simulate five scRNA-seq datasets which have different size and different sparsity for more comprehensive analysis. We set *group.prob* to (0.65, 0.25, 0.1) for all simulated datasets, and change the scale and sparsity by adjusting *nCells* and *dropout.mid* respectively. The other parameters are set to default. The samples of the five simulated datasets are 1000, 1000, 1000, 500, 1500, and the corresponding sparsity is 0.61, 0.8, 0.94, 0.94, 0.94, respectively.

**Table 1.** The details of real scRNA-seq datasets used in this study.

| Dataset | No. of cells | No. of genes | No. of groups | Units |
| --- | --- | --- | --- | --- |
| Treutlein [32] | 80 | 959 | 5 | FPKM |
| Yan [33] | 90 | 20,214 | 7 | RPKM |
| Deng [34] | 135 | 12,548 | 7 | RPKM |
| Goolam [35] | 124 | 40,315 | 5 | CPM |
| Ting [36] | 114 | 14,405 | 5 | RPM |
| Song [37] | 214 | 27,473 | 4 | TPM |
| Engel [38] | 203 | 23,337 | 4 | TPM |
| Haber [39] | 1522 | 20,108 | 9 | TPM |
| Vento [40] | 5418 | 33,693 | 38 | HTSeq-count |
| Macosko [41] | 6418 | 12,822 | 39 | UMI |

*Note:* FPKM, fragments per kilobase of exon model per million mapped fragments; RPKM, reads per kilobase of exon model per million mapped reads; CPM/RPM, counts /reads of exon model per million mapped reads; TPM, transcripts per kilobase of exon model per million mapped reads; UMI, unique molecular identifiers.

### scRNA-seq clustering methods

For performance comparison, we take the original SSR and eight state-of-the-art clustering methods, *i.e.* SIMLR [8], MPSSC [19], Corr [11], SNN-Cliq [10], NMF [13], SC3 [15], dropClust [43], and Seurat [44] as comparison. Among these methods, SIMLR, MPSSC, Corr, and SNN-Clip focus on similarity learning. Both SIMLR and MPSSC learn a representative similarity matrix from multi-Gaussian-kernels with different resolutions. Corr introduces a cell-pair differentiability correlation to relieve the effect of drop-outs. SNN-Cliq applies the shared-nearest-neighbor to redefine the pairwise similarity. NMF detects the type of cells by projecting the high dimensional data into a latent space, in which each dimension of the latent space denotes a specific type. SC3 is a typical and powerful consensus clustering method. It obtains clusters by applying different upstream processes and the final clusters are desired to fit better. DropClust is a clustering algorithm designed for large-scale single cell data, and it exploits an approximate nearest neighbour search technique to reduce the time complexity of analyzing large-scale data. Seurat, a popular R package for single cell data analysis, obtains cell groups based on KNN-graph and Louvain clustering. Moreover, the native spectral clustering [29] with the Pearson similarity is considered as a baseline.

### Metric of performance evaluation

We evaluate the proposed approach using two common metrics, *i.e.* normalized mutual information (NMI) [45] and adjusted rand index (ARI) [46] which have been widely used to assess clustering performance. Both NMI and ARI evaluate the consistency between the obtained clustering and pre-annotated labels while have a slightly different on the emphases [47]. Given the real labels *L*1 and the clustering labels *L*2, NMI is calculate as

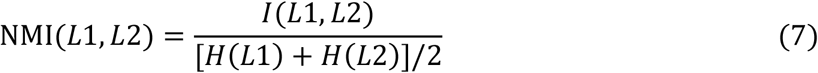

*I* (*L*1, *L*2) is the mutual information between *L*1 and *L*2 and *H* denotes entropy. For ARI, given *L*1 and *L*2, it is computed as

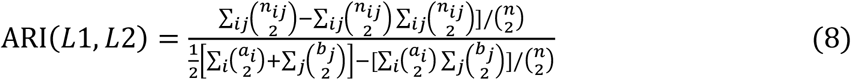

where *n*_*ij*_ is the number of cells in both group *L*1_*i*_ and group *L*2_*j*_, *a*_*i*_ and *b*_*j*_ denote the number of cells in group *L*1_*i*_ and group *L*2_*j*_ respectively.

## Results and discussion

### Cell type identification and comparative analysis

In order to evaluate the performance of SSRE comprehensively, we first apply it on ten pre-annotated real scRNA-seq datasets and compare its performance with the original SSR, the native SC and eight state-of-the-art clustering methods from different categories. See details in the Materials and methods section. Then, we perform all these methods on five simulated datasets for further comparison. In our experiments, for a fairer comparison, we set the number of clusters of all methods to the number of pre-annotated types for all methods except SNN-Cliq and Seurat, because SNN-Cliq and Seurat does not need the number of clusters as input. The other parameters in all the methods are set to the default values described in the original papers. **Table 2** and **Table 3** summarizes the NMI and ARI values of all methods on ten real scRNA-seq datasets respectively. The results of Corr in large datasets are unreachable because of the high computational complexity. As shown in Table 2 and Table 3, the proposed method SSRE outperforms all other methods in most cases. SSRE achieves the best or tied first on seven datasets upon NMI and ARI. Moreover, SSRE ranks second on three datasets based on NMI and two datasets based on ARI respectively. It demonstrates that SSRE obtains more reliable results independent to the scale and the biological conditions of scRNA-seq data. When is compared with original SSR, SSRE performs better in all of the datasets regarding NMI and ARI, which validates the effectiveness of the enhancement strategy in SSRE. The results of simulation experiment are shown in Table S1 and Table S2. We can see that SSRE has the better performance overall in terms of NMI and ARI, which shows the good stability of SSRE under different conditions. SSRE is slightly time-consuming compared with some methods like SC, Seurat, and dropClust, but is still in the reasonable range. More detailed descriptions can be found in section 2 of File S1.

**Table 2.** NMI values of all analyzed methods across ten real datasets.

| Methods | Treutlein | Yan | Deng | Goolam | Ting | Song | Engel | Haber | Vento | Macosko |
| --- | --- | --- | --- | --- | --- | --- | --- | --- | --- | --- |
| SC | 0.71 | 0.69 | 0.63 | 0.72 | 0.89 | 0.51 | 0.71 | 0.40 | 0.70 | 0.80 |
| SNN-Cliq | 0.64 | 0.76 | 0.78 | 0.62 | 0.73 | 0.54 | 0.31 | 0.24 | 0.51 | 0.55 |
| SIMLR | 0.69 | 0.79 | **0.84** | 0.56 | 0.98 | 0.67 | 0.74 | 0.40 | 0.64 | 0.72 |
| SC3 | 0.73 | 0.81 | 0.72 | 0.72 | **1.00** | **0.73** | **0.81** | 0.05 | 0.66 | 0.83 |
| NMF | 0.67 | 0.64 | 0.68 | 0.55 | 0.60 | 0.52 | 0.70 | 0.05 | 0.68 | 0.72 |
| MPSSC | 0.80 | 0.76 | 0.76 | 0.56 | 0.98 | 0.60 | 0.55 | 0.17 | 0.40 | 0.71 |
| Corr | 0.64 | 0.81 | 0.72 | 0.56 | 0.71 | 0.60 | 0.29 | - | - | - |
| dropClust | **0.82** | 0.76 | 0.73 | 0.81 | 0.91 | 0.61 | 0.29 | 0.43 | 0.67 | 0.71 |
| Seurat | 0.53 | 0.72 | 0.68 | 0.62 | 0.80 | 0.71 | 0.72 | **0.62** | 0.69 | 0.62 |
| SSR | 0.73 | 0.86 | 0.79 | 0.69 | **1.00** | 0.69 | 0.76 | 0.52 | 0.70 | 0.84 |
| SSRE | **0.82** | **0.92** | 0.81 | **0.83** | **1.00** | **0.73** | 0.77 | 0.53 | **0.72** | **0.87** |

**Table 3.** ARI values of all analyzed methods across ten real datasets.

| Methods | Treutlein | Yan | Deng | Goolam | Ting | Song | Engel | Haber | Vento | Macosko |
| --- | --- | --- | --- | --- | --- | --- | --- | --- | --- | --- |
| SC | 0.59 | 0.44 | 0.33 | 0.54 | 0.89 | 0.49 | 0.67 | 0.19 | 0.37 | 0.52 |
| SNN-Cliq | 0.26 | 0.49 | 0.54 | 0.20 | 0.55 | 0.27 | 0.13 | 0.00 | 0.03 | 0.07 |
| SIMLR | 0.51 | 0.60 | **0.67** | 0.30 | 0.98 | 0.55 | 0.67 | 0.21 | 0.38 | 0.52 |
| SC3 | 0.65 | 0.71 | 0.47 | 0.54 | **1.00** | 0.70 | 0.71 | 0.09 | 0.40 | 0.77 |
| NMF | 0.47 | 0.42 | 0.44 | 0.30 | 0.29 | 0.31 | 0.62 | 0.06 | 0.45 | 0.51 |
| MPSSC | 0.61 | 0.60 | 0.48 | 0.40 | 0.98 | 0.50 | 0.48 | 0.10 | 0.16 | 0.43 |
| Corr | 0.56 | 0.71 | 0.53 | 0.32 | 0.50 | 0.41 | 0.13 | - | - | - |
| dropClust | **0.88** | 0.62 | 0.46 | 0.59 | 0.89 | 0.58 | 0.24 | 0.24 | 0.45 | 0.45 |
| Seurat | 0.57 | 0.64 | 0.53 | 0.53 | 0.73 | 0.66 | 0.69 | **0.43** | 0.46 | 0.33 |
| SSR | 0.51 | 0.79 | 0.56 | 0.49 | **1.00** | 0.63 | 0.74 | 0.31 | 0.45 | 0.73 |
| SSRE | 0.62 | **0.91** | 0.65 | **0.67** | **1.00** | **0.75** | **0.75** | 0.32 | **0.47** | **0.86** |

Estimating number of clusters is another key step in most clustering methods, which affects the accuracy of clustering method. In SSRE, we perform eigengap [29] on the learned similarity matrix to estimate the number of clusters. Eigengap is a typical cluster number estimation method, and it determines the number of clusters by calculating max gap between eigenvalues of a Laplacian matrix. To assess reliability of the estimation in different methods, we compare their estimated numbers and pre-annotated number. The results are summarized in Table S3. Besides SSRE and SSR, another four methods which also focus on similarity learning are selected for comparison. More experimental details can be seen in section 3 of File S1.

### Parameter selection and analysis

In SSRE, four parameters are required to be set by users, *i.e.* penalty coefficients *λ* and *γ* in solving sparse similarity *sim*_*sparse*_, gene selection threshold *t*, and the number of nearest neighbors *k* in similarity enhancement procedure. The selection of the threshold *t* can be determined adaptively by solving Equation (4) described in Section data preprocessing and gene selection. For the number of nearest neighbors *k*, we set *k* = 0.1 ∗ *n* (*n* is the number of cells) as default in small datasets with less than 5000 cells and *k* = 100 in other larger datasets. The other two parameters *λ* and *γ* in augmented Lagrangian (we use 1/*λ* and 1/*γ* in the coding implementation) are proportionally set as

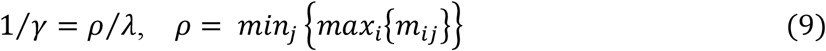

where *m*_*ij*_ is the element of matrix *M* = *X*^*T*^*X* and it is equivalent to the cosine similarity between cells *x*_*i*_ and *x*_*j*_, which is the same as previous work [18]. In our experiments, *ρ*/*λ* is set to a constant. So, for given dataset, the larger value of *ρ* will lead to the larger value of *λ*, which will result in the sparser matrix C. It means that the value of *ρ* can control the sparsity of matrix C adaptively in different datasets. Moreover, to validate the effect of penalty coefficient *λ* in clustering results, we test our model with *ρ*/*λ* from 2 to 30 with the increment of 2 on all real datasets. As shown in **Figure 2**, the corresponding ARI and NMI indicate that the performance of SSRE is basically stable when *ρ*/*λ* is in the interval of 6 and 20 (the resting results are shown in Supplementary Figure S1). In our study, we set *ρ*/*λ* to 10 and 1/*λ* = *ρ*/*λ* as default for all datasets.

**Figure 2.**
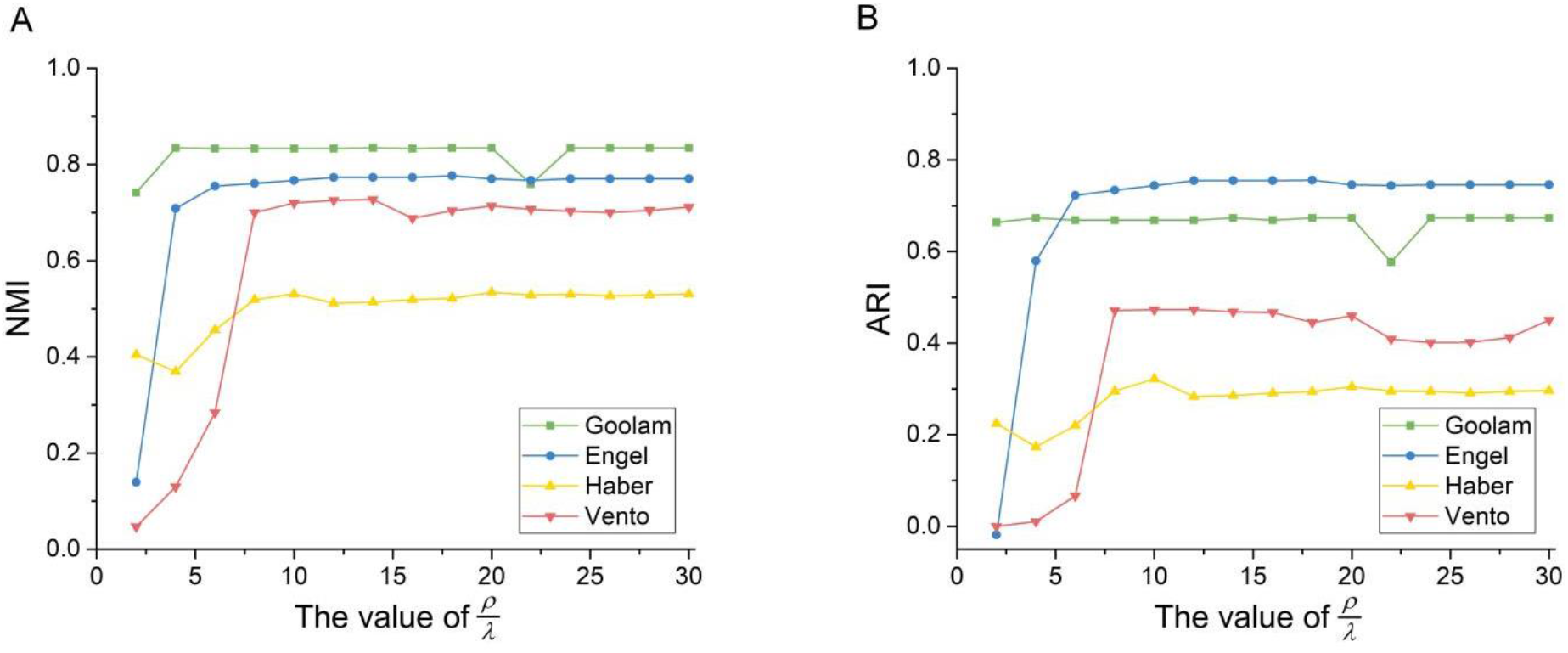
Analysis of parameter setting in SSRE. **A.** NMI values of SSRE on the Goolam, Engel, Haber, Vento datasets with different *ρ*/*λ*. **B.** ARI values of SSRE on the Goolam, Engel, Haber, Vento datasets with different *ρ*/*λ*.

### Visualization

One of the most valuable aims in single cell analysis is to identify new cell types or subtypes [6]. Visualization is an effective tool to give an intuitive display of the subgroups in all cells. The t-distributed stochastic neighbor embedding (t-SNE) [48] is one of the most popular visualization methods and has been proved powerful in scRNA-seq data. In this section, we perform a modified t-SNE on learned similarities to project high dimensional data into two-dimensional space. We focus on two datasets Goolam and Yan and select the native t-SNE, SIMLR, MPSSC, Corr based t-SNE for comparison. In Goolam [35], cells are derived from mouse embryos in five differentiation stages: 2-cell, 4-cell, 8-cell, 16-cell and 32-cell. Taking learned similarities of Goolam as input, the visualization results are shown in **Figure 3 (A, B, C, D, E, F)**. SSRE places cells with the same type together and distinguishes cells with different types clearly. The groups in SIMLR are clearly distinguished from each other but some cells with the same type are separated. The second dataset Yan [33] is obtained from human pre-implantation embryos and involves seven primary stages of preimplantation development: metaphase II oocyte, zygote, 2-cell, 4-cell, 8-cell, morula and late blastocyst. **Figure 3 (G, H, I, J, K, L)** shows the results of Yan dataset. We can see that Corr, SIMLR, and SSRE have a better overall performance than other methods. However, the four cell types, *i.e.*, oocyte, zygote, 2-cell, and 4-cell, are mixed totally in Corr and partially in SIMLR. Moreover, SIMLR also divides the cells with the same type into distinct groups which are usually far away from each other. SSRE groups cells more accurately, according to oocyte, 2-cell, and other cells than the competing methods.

**Figure 3.**
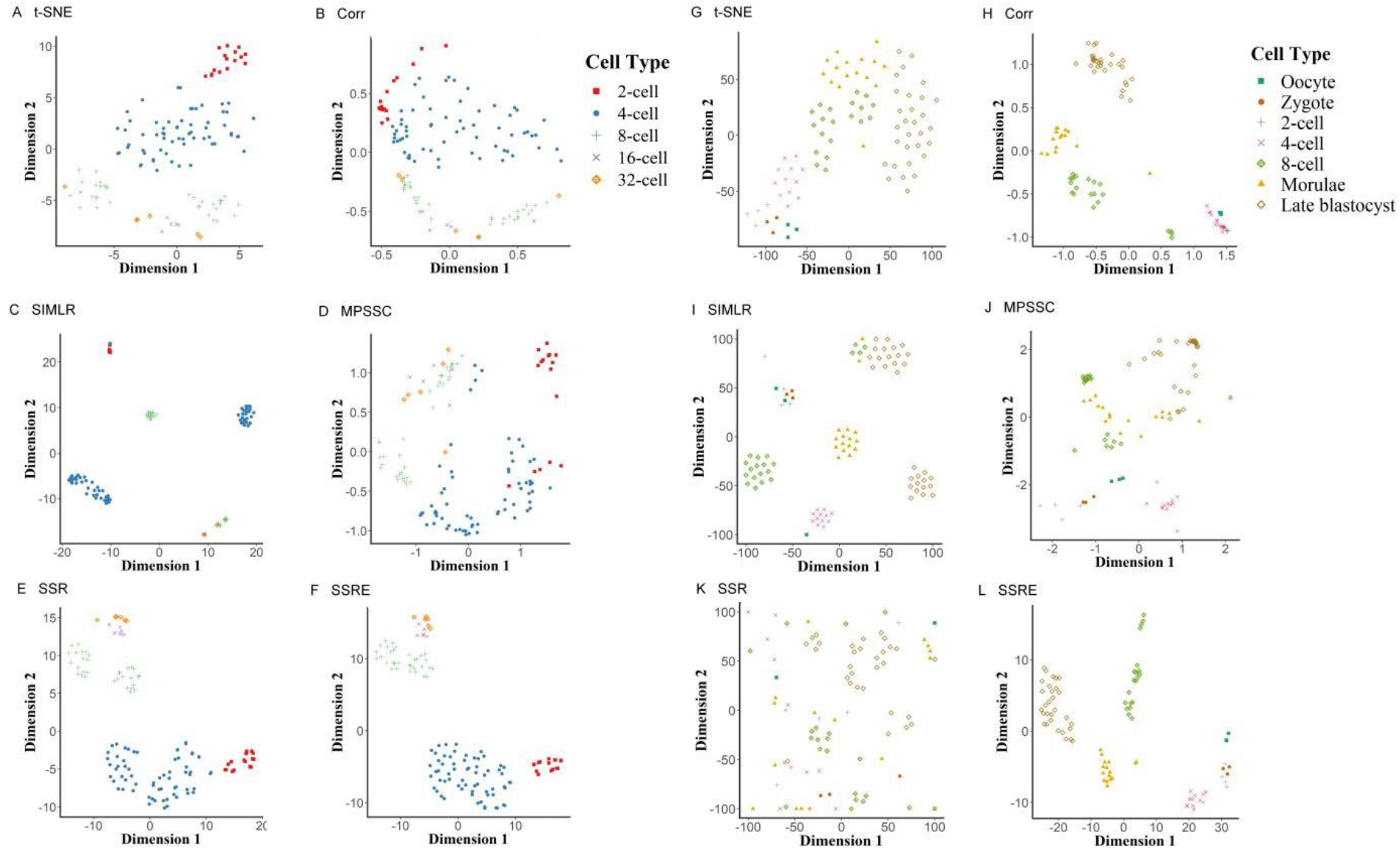
Visualization of cells by different methods. The 2D visualization of the cells in Goolam dataset by using t-SNE (A), Corr (B), SIMLR (C), MPSSC (D), SSR (E), SSRE (F), and in Yan dataset by using the same six methods, t-SNE (G), Corr (H), SIMLR (I), MPSSC (J), SSR (K), and SSRE (L).

### Identification of differentially expressed genes

The predicted clusters may potentially enable enhanced downstream scRNA-seq data analysis in biological sights. As a demonstration, here we aim to detect significantly differentially expressed genes based on the clustering results. Specifically, we apply the Kruskal-Wallis test [49] to the gene expression profiles with the inferred labels. The Kruskal-Wallis test, a non-parametric method, is often used for testing if two or more groups are from the same distribution. We use the R function *kruskal.test* to perform the Kruskal-Wallis test and calculate differential expression according to the P-value. The significant P-value (P < 0.01) of a gene indicates that the gene’s expression in at least one group stochastically dominates one other group. We use the Yan [33] dataset as an example to analyze the differential expressed genes. The details of Yan have been introduced above. Supplementary Figure S2 shows the heat map of gene expression of the detected 50 most significantly differentially expressed genes. Notice that genes *NLRP11*, *NLRP4*, *CLEC10A*, *H1FOO*, *GDF9*, *OTX2*, *ACCSL*, *TUBB8*, and *TUBB4Q* have been reported in previous studies [33,50] and are also identified by SSRE. Genes *CLEC10A*, *H1FOO*, and *ACCSL* are reported as the markers of 1-cell stage cells (Zygote) of human early embryos while *NLRP11* and *TUBB4Q* are the markers of 4-cell [51]. Genes *GDF9* and *OTX2* are the markers of germ cell and primitive endoderm cell, respectively [52,53]. Genes *H1FOO* and *GDF9* are marked as the potential stage-specific genes in the oocyte and the blastomere of 4-cell stage embryos [54]. Certain *PRAMEF* family genes are reported as ones with transiently enhanced transcription activity in 8-cell stage. *MBD3L* family genes are identified as 8-cell-genes during the human embryo development in the previous studies [55,56]. All these are part of the most 50 significantly differentially expressed genes detected by SSRE.

## Conclusion

Identifying cell types from single cell transcriptome data is a meaningful but challengeable work because of the high-level noise and high dimension. The ideal identification of cell types enables more reliable characterizations of a biological process or phenomenon, otherwise introducing even more biases. Many approaches from different perspectives have been proposed recently, but the accuracy of cell type identification is still far from expectation. In this paper, we proposed SSRE, a computational framework focused on similarity learning, for cell type identification and visualization of scRNA-seq data. Besides three classical pairwise similarities, SSRE computes the sparse representation similarity of cells based on the subspace theory. Moreover, we designed a gene selection process and an enhancement strategy based on the characteristics of different similarities to learn more reliable similarities. We expect that by appropriately combining multiple similarity measures and adopting the embedding of sparse structure, SSRE can further improve the clustering performance. With systematic performance evaluation on multiple scRNA-seq datasets, it shows that SSRE achieves superior performance among all competing methods. Furthermore, the further downstream analyses demonstrate that the learned similarity and inferred clusters can potentially be applied on more exploratory analysis, such as identifying gene markers, detecting new cell subtypes and so on. In addition, for a more flexible usage, in our implementation code, users can choose one or two of three correlation similarities mentioned in this study to perform gene selection and similarity enhancement procedure, and the default is all three correlation similarities. Nonetheless, the proposed computational framework allows some future improvements. For instance, selecting gene sets and combining similarities by considering multiple factors simultaneously [57,58], integrating multi-omics data [59,60] for similarity learning, and using parallel computing in clustering [61] to reduce time consume.

## Data availability

The real scRNA-seq datasets used in this paper can be obtained from GEO (Treutlein: <u>GSE52583</u>, Yan: <u>GSE36552</u>, Deng: <u>GSE45719</u>, Ting: <u>GSE51372</u>, <u>GSE60407</u>, and <u>GSE51827</u>, Song: <u>GSE85908</u>, Engel: <u>GSE74597</u>, Haber: <u>GSE92332</u>, and Macosko: <u>GSE63473</u>) and ArrayExpress (Goolam: <u>E-MTAB-3321</u>, Vento: <u>E-MTAB-6678</u>).

## Authors’ contributions

ZL and ML conceived and designed the experiments. ZL, XY wrote and revised the code. ZL, YT, RZ performed the experiments and analyzed the data. ZL, RZ and ML drafted the manuscript. JC, FXW, JW revised the manuscript. All authors read and approved the final manuscript.

## Acknowledgements

This work was supported in part by the NSFC-Zhejiang Joint Fund for the Integration of Industrialization and Information under Grant No. U1909208; the 111 Project (No. B18059); the Hunan Provincial Science and Technology Program (2018WK4001); the Fundamental Research Funds for the Central Universities-Freedom Explore Program of Central South University (No. 2019zzts592); and US National Natural Science Foundation (no.1716340).

