## Supplementary File, Figures and Tables for "SSRE: Cell Type Detection Based on Sparse Subspace Representation and Similarity Enhancement"

### Supplementary materials for "SSRE: cell type detection based on sparse subspace representation and similarity enhancement"

Zhenlan Liang[1], Min Li[1,*], Ruiqing Zheng[1], Yu Tian[1], Xuhua Yan[1], Jin Chen[2], Fang-Xiang Wu[3], Jianxin Wang[1]

*[1] School of Computer Science and Engineering, Central South University, Changsha 410083, China*

*[2] College of Medicine, University of Kentucky, Lexington 40536, USA*

*[3] Division of Biomedical Engineering, University of Saskatchewan, Saskatoon SKS7N5A9, Canada*

* Corresponding author

 (Li M)

#### Section 1: The ADMM used in SSRE

According to the optimization program (2) in the main context, the augmented Lagrangian formulation is written as follows:

$$\mathcal{L}_{\frac{1}{\gamma}}(Z,C,Y) = \frac{1}{2\lambda}\|X - XZ\|_F^2 + \|C\|_1 + tr(Y^T(Z - C)) + \frac{1}{2\gamma}\|C - Z\|^2 \qquad (S1)$$

where *X* is the gene expression matrix, *C* is the target sparse representation matrix to be solved, *J* is an auxiliary matrix, *Y* is the dual variable or Lagrange multiplier and $\gamma$ is a user-defined parameter. The ADMM updates the one of matrix *C*, *Z*, *Y* by fixing others each time with the following formulas

$$Z^{k+1} = \text{argmin}_Z\ \mathcal{L}_{\frac{1}{\gamma}}(Z, C^k, Y^k) \qquad (S2)$$

$$= \text{argmin}_Z \left\{\frac{1}{2\lambda}\|X - XZ\|_F^2 + \|C^k\|_1 + tr\left(Y^{k^T}(Z - C^k)\right) + \frac{1}{2\gamma}\|C^k - Z\|^2\right\}$$

$$= \left(\frac{X^TX}{\lambda} + \frac{1}{\gamma}\right)^{-1}\left(\frac{X^TX}{\lambda} - Y^{k^T} + \frac{1}{\gamma}C^k\right)$$

$$C^{k+1} = \text{argmin}_C\ \mathcal{L}_{\frac{1}{\gamma}}(Z^{k+1}, C, Y^k) \qquad (S3)$$

$$= \text{argmin}_C \left\{\frac{1}{2\lambda}\|X - XZ^{k+1}\|_F^2 + \|C\|_1 + tr\left(Y^{k^T}(Z^{k+1} - C)\right) + \frac{1}{2\gamma}\|C - Z^{k+1}\|^2\right\}$$

$$= Soft_{\lambda,\gamma}\left(Z^{k+1} + \gamma Y^k\right)$$

$$Y^{k+1} = Y^k + \frac{1}{\gamma}(Z^{k+1} - C^{k+1}) \qquad (S4)$$

where $Soft_{\lambda,\gamma}(*)$ is a soft-thresholding operator [1].

#### Section 2: Results on simulated datasets

Besides real datasets, we also apply SSRE on five simulated datasets with different size or sparsity. The size and sparsity of five simulated datasets are as: Sim_data_1 (size: 1000cells, sparsity:0.61), Sim_data_2 (size: 1000cells, sparsity:0.8), Sim_data_3 (size: 1000cells, sparsity:0.94), Sim_data_4 (size: 500cells, sparsity: 0.94), Sim_data_5 (size: 1500cells, sparsity:0.94). Table S1 and Table S2 summarize the results on datasets with different sparsity and different size respectively, where the results of Corr in Sim_data_5 is unreachable because of high computational complexity. We can see that SSRE has the best performance overall in two simulation experiments in terms of NMI and ARI. DropClust gets the smallest running time on almost all datasets, but the running time needed by SSRE is also acceptable

**Table S1 Results of all analyzed methods on simulated datasets with different sparsity**

| **Methods** | **Sim_data_1** | | | **Sim_data_2** | | | **Sim_data_3** | | |
|---|---|---|---|---|---|---|---|---|---|
| | **NMI** | **ARI** | **Time(s)** | **NMI** | **ARI** | **Time(s)** | **NMI** | **ARI** | **Time(s)** |
| SC | 0.98 | 0.99 | 8.79 | 0.95 | 0.97 | 7.68 | 0.61 | 0.63 | 7.34 |
| SNN-Cliq | 0.43 | 0.03 | 24.21 | 0.42 | 0.03 | 25.58 | 0.28 | 0.01 | 25.54 |
| SIMLR | 0.98 | 0.99 | 49.76 | 0.97 | 0.99 | 48.70 | 0.42 | 0.35 | 32.55 |
| SC3 | **1.00** | **1.00** | 1404.4 | **0.99** | **1.00** | 1323.1 | 0.67 | 0.69 | 1393.0 |
| NMF | 0.96 | 0.98 | 403.28 | 0.93 | 0.96 | 183.27 | 0.62 | 0.67 | 55.03 |
| MPSSC | 0.48 | 0.41 | 29.63 | 0.53 | 0.43 | 84.31 | 0.39 | 0.36 | 26.87 |
| Corr | 0.64 | 0.75 | 37,144 | 0.59 | 0.70 | 35,300 | 0.02 | -0.01 | 34,302 |
| dropClust | 0.80 | 0.65 | **8.78** | 0.76 | 0.57 | **6.90** | 0.45 | 0.41 | **6.14** |
| Seurat | 0.73 | 0.52 | 18.03 | 0.81 | 0.61 | 16.95 | 0.66 | 0.55 | 15.58 |
| SSR | **1.00** | **1.00** | 35.56 | **0.99** | **1.00** | 43.92 | 0.65 | 0.72 | 46.26 |
| SSRE | **1.00** | **1.00** | 65.99 | **0.99** | 0.99 | 59.44 | **0.69** | **0.80** | 56.32 |

**Table S2 Results of all analyzed methods on simulated datasets with different size**

| **Methods** | **Sim_data_4** | | | **Sim_data_3** | | | **Sim_data_5** | | |
|---|---|---|---|---|---|---|---|---|---|
| | **NMI** | **ARI** | **Time(s)** | **NMI** | **ARI** | **Time(s)** | **NMI** | **ARI** | **Time(s)** |
| SC | 0.68 | 0.71 | **2.98** | 0.61 | 0.63 | 7.34 | 0.38 | 0.37 | 12.50 |
| SNN-Cliq | 0.35 | 0.02 | 7.00 | 0.28 | 0.01 | 25.54 | 0.22 | 0.01 | 61.63 |
| SIMLR | 0.46 | 0.38 | 12.66 | 0.42 | 0.35 | 32.55 | 0.31 | 0.30 | 78.93 |
| SC3 | **0.95** | **0.98** | 858.5 | 0.67 | 0.69 | 1393.0 | 0.42 | 0.54 | 3075.8 |
| NMF | 0.78 | 0.83 | 43.25 | 0.62 | 0.67 | 55.03 | 0.46 | 0.54 | 73.03 |
| MPSSC | 0.57 | 0.43 | 5.10 | 0.39 | 0.36 | 26.87 | 0.44 | 0.37 | 338.10 |
| Corr | 0.02 | 0.01 | 5123.10 | 0.02 | -0.01 | 34,302 | - | - | - |
| dropClust | 0.64 | 0.50 | 3.18 | 0.45 | 0.41 | **6.14** | 0.31 | 0.24 | **8.84** |
| Seurat | 0.68 | 0.57 | 14.83 | 0.66 | 0.55 | 15.58 | **0.76** | 0.63 | 18.64 |
| SSR | 0.92 | 0.96 | 11.60 | 0.65 | 0.72 | 46.26 | 0.57 | 0.65 | 79.78 |
| SSRE | **0.95** | **0.98** | 13.92 | **0.69** | **0.80** | 56.32 | 0.63 | **0.72** | 103.36 |

#### Section 3: The estimation of number of clusters

With the learned similarity of SSR and SSRE, we apply the eigengap [2] to estimate the number of clusters. Moreover, we choose SIMLR [3], MPSSC [4], Corr [5], and SNN-Cliq [6] as the competing methods which also focus on similarity learning. For these compared methods, we perform the corresponding estimation algorithm provided by themselves. Table S3 summarizes the results on ten real datasets. It shows that the proposed method SSRE gets the same number as pre-annotated number in two datasets (Ting [7] and Vento [8]) and closest with pre-annotated number in most datasets (Treutlein [9], Goolam [11], Song [12], Haber [13] and Macosko [14]). We can see that there is no method can estimate the number of clusters as same as pre-annotated numbers in all datasets, and SSRE has a better performance overall.

**Table S3 The number of clusters estimated by different methods**

| Dataset | pre-annotated number | SSRE | SSR | SNN-Cliq | Corr | SIMLR | MPSSC |
|---|---|---|---|---|---|---|---|
| Treutlein [9] | 5 | 4 | 4 | 14 | 3 | 10 | 15 |
| Yan [15] | 7 | 14 | 16 | 18 | 5 | 12 | 16 |
| Deng [10] | 7 | 5 | 5 | 8 | 2 | 9 | 17 |
| Goolam [11] | 5 | 3 | 8 | 17 | 3 | 11 | 15 |
| Ting [7] | 5 | 5 | 5 | 8 | 3 | 5 | 15 |
| Song [12] | 4 | 3 | 4 | 30 | 2 | 3 | 14 |
| Engel [16] | 4 | 1 | 1 | 13 | 2 | 3 | 2 |
| Haber [13] | 9 | 8 | 15 | 301 | - | 10 | 15 |
| Vento [8] | 38 | 38 | 40 | 178 | - | 11 | 1 |
| Macosko [14] | 39 | 42 | 49 | 548 | - | 6 | 11 |

#### Section 4: Results with different values of $\rho/\lambda$ on more real datasets

To validate the effect of user-defined penalty coefficient $\boldsymbol{\lambda}$ (we use $\mathbf{1}/\boldsymbol{\lambda}$ in matlab code), we set $\boldsymbol{\rho}/\boldsymbol{\lambda}$ from 2 to 30 with the increment of 2. In addition to the four real datasets in the main context, the corresponding ARI and NMI of other real datasets are shown in Figure S1.

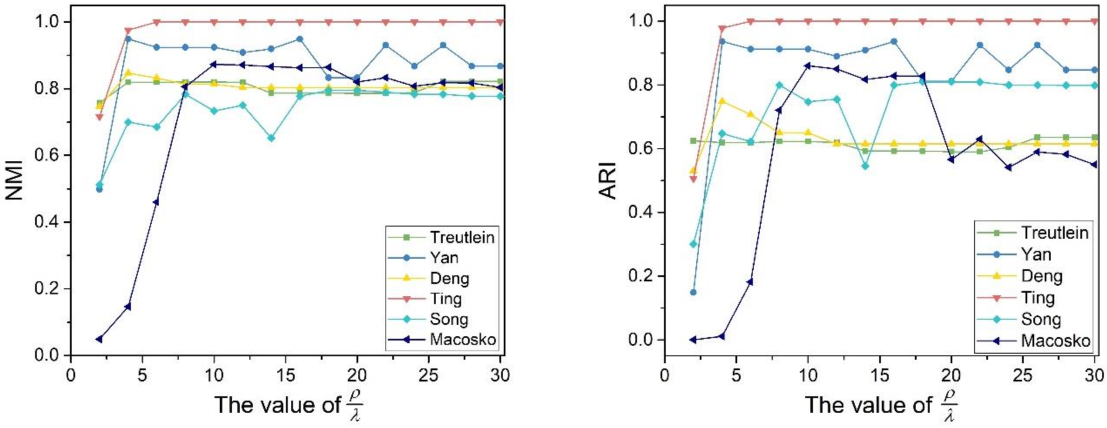


**Figure S1 Analysis of parameter setting in SSRE**

**A.** NMI values of SSRE on six real scRNA-seq datasets with different $\rho/\lambda$. **B.** ARI values of SSRE on the six datasets with different $\rho/\lambda$.

#### Section 5: Identification of differentially expressed genes

With the predicted clusters by SSRE, we try to detect significant differentially expressed genes based on Kruskal-Wallis test. The heat map of gene expression of the top 50 most significant differentially expressed genes in Yan is displayed in Figure S2.

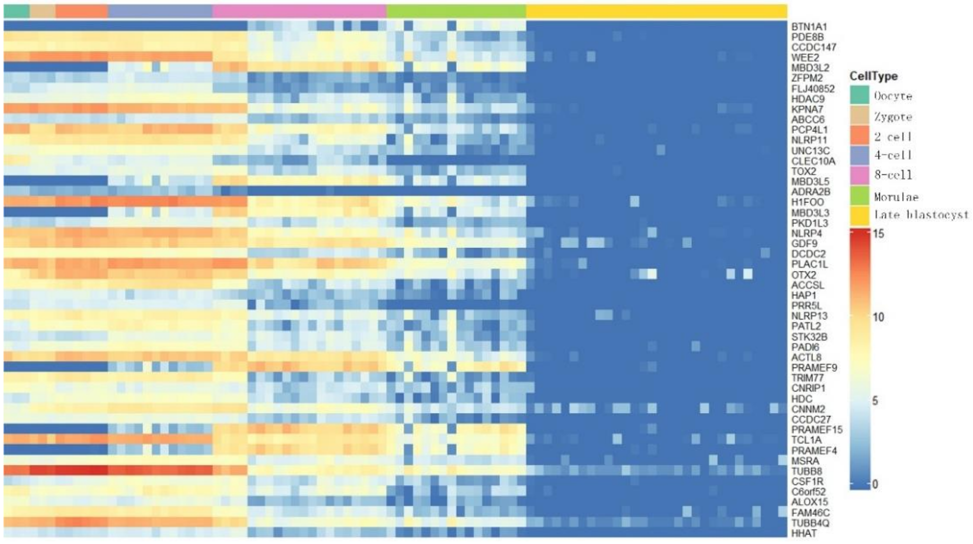


**Figure S2 Analysis of differentially expressed genes**

The heat map of top 50 most significant differentially expressed genes identified by SSRE on Yan dataset.
